# Demographic History Inference and the Polyploid Continuum

**DOI:** 10.1101/2022.09.15.508148

**Authors:** Paul D. Blischak, Mathews Sajan, Michael S. Barker, Ryan N. Gutenkunst

**Author notes:** Corresponding authors: Department of Molecular and Cellular Biology, University of Arizona, Tucson, AZ 85721, USA.;.

## Abstract

Polyploidy is an important generator of evolutionary novelty across diverse groups in the Tree of Life, including many crops. However, the impact of whole-genome duplication (WGD) depends on the mode of formation: doubling within a single lineage (autopolyploidy) versus doubling after hybridization between two different lineages (allopolyploidy). Researchers have historically treated these two scenarios as completely separate cases based on patterns of chromosome pairing, but these cases represent ideals on a continuum of chromosomal interactions among duplicated genomes. Understanding the history of polyploid species thus demands quantitative inferences of demographic history and rates of exchange between subgenomes. To meet this need, we developed diffusion models for genetic variation in polyploids with subgenomes that cannot be bioinformatically separated and with potentially variable inheritance patterns, implementing them in the dadi software. We validated our models using forward SLiM simulations and found that our inference approach is able to accurately infer evolutionary parameters (timing, bottleneck size) involved with the formation of auto- and allotetraploids, as well as exchange rates in segmental allotetraploids. We then applied our models to empirical data for allotetraploid shepherd’s purse (*Capsella bursa-pastoris*), finding evidence for allelic exchange between the subgenomes. Taken together, our model provides a foundation for demographic modeling in polyploids using diffusion equations, which will help increase our understanding of the impact of demography and selection in polyploid lineages.

---

Polyploidy, or whole genome duplication (WGD), is a mechanism for potentially rapid evolutionary change. Many lineages in the Tree of Life have experienced WGD events in their ancient pasts (“paleopolyploids”; Ohno 1970; Furlong and Holland 2001; Cui et al. 2006; Jiao et al. 2011; Li et al. 2018; Leebens-Mack et al. 2019; Li and Barker 2020) and the formation of recent polyploids (“neopolyploids”) is especially common in plants and some groups of animals (e.g., amphibians Otto and Whitton 2000; Gregory and Mable 2005; Wood et al. 2009). The prevalence of polyploidy events through deep time as well as in the present have led many to hypothesize that polyploids are able to tolerate different or more extreme environments/conditions than their diploid progenitors (Comai 2005; Baduel et al. 2018; Baniaga et al. 2020; Van de Peer et al. 2020). And although selection and adaptation has been quantified in some studies of polyploids (Selmecki et al. 2015; Arnold et al. 2016; McIntyre and Strauss 2017; Monnahan et al. 2019), there remains a lack of consensus surrounding the role of demographic versus selective processes in contributing to the evolutionary trajectories of polyploids after their formation (Blischak et al. 2018b; Li et al. 2021).

One reason for this lack of consensus is the difficulty of building demographic models that accommodate the additional set(s) of chromosomes that polyploids possess plus their potential interactions. The mode of formation for a polyploid, either through WGD within a lineage (autopolyploidy) or following hybridization between two or more different lineages (allopolyploidy), has a great impact on how the lineage evolves post-WGD. These differences result from the patterns of chromosomal interactions that occur in autopolyploids versus allopolyploids, with autopolyploids ranging from free recombination among all chromosomes in the genome to allopolyploids with only recombination between chromosomes from the same parental lineage. Previous work on the demography of polyploids has typically assumed that the species under study falls into one of these two categories. However, the existence of intermediate types of polyploids (“segmental allopolyploids”; Stebbins 1950) challenges the placement of polyploids into these discrete categories, suggesting that polyploids may be better described by a continuum (Gaut and Doebley 1997; Meirmans and van Tienderen 2013; Mason and Wendel 2020). Building this continuum-like nature into a demographic model has not been explored extensively other than in a few case studies. For example, Roux and Pannell (2015) and Roux *et al*. (2021) used approximate Bayesian computation to infer the mode of origin of a polyploid by simulating data from different types of polyploids, using migration as a means to simulate polyploid data showing a mixed pattern of allelic inheritance, and comparing the simulated genetic data to the empirical data using summary statistics. Approaches such as this offer a promising avenue for further development of more generalized demographic models for different types of polyploids, but are also limited by their reliance on simulations and the comparison of summary statistics rather than on likelihood-based parameter estimation and model comparison.

Another important issue for polyploids is the additional complexity of standard bioinformatic procedures, such as read mapping and variant calling, in the presence of duplicated chromosomes. Because the high-throughput sequencing reads collected for reference genome-based SNP calling are often short (*∼* 50– 250 bp depending on the platform), reads can potentially map to multiple chromosomes, leading either to large amounts of discarded data if only uniquely mapped reads are kept or to the identification of erroneous SNPs due to read mismapping. One way to help alleviate this issue is to focus on calling SNPs at the correct ploidy level of the sequenced individual, rather than trying to ensure that reads are mapping to the appropriate subgenome and calling SNPs separately at the ploidy level of each subgenome (e.g., calling tetraploid genotypes instead of calling genotypes in two diploid subgenomes). For autopolyploids, this method of SNP calling is simpler since their homoeologous chromosomes are derived from the same lineage and several methods for genotyping in this scenario exist (Serang et al. 2012; Blischak et al. 2018a; Gerard et al. 2018; Clark *et al*. 2019). However, for allopolyploid and segmental allopolyploid lineages, divergence between the parental subgenomes means that identifying homoeologous positions for genotyping is more difficult, and it is further complicated by having to distinguish between SNPs within a subgenome and fixed differences between subgenomes. Models to separately estimate SNPs within allopolyploid subgenomes have been proposed and used in a variety of crop species (Blischak et al. 2018a; Clevenger and Ozias-Akins 2015; Clevenger et al. 2018; Korani et al. 2019; Clark *et al*. 2019; Kulkarni et al. 2022), but they typically require knowledge about all parental subgenomes. If the nature of a polyploid’s formation mode is unknown or if parental information is not available for an allopolyploid, then these approaches are not able to be used. One possible solution would be to simply genotype a polyploid at its ploidy level without any attempt to separate out lower-ploidy subgenomes. With this approach, the task of determining the mode of polyploid formation could be incorporated into any downstream analyses rather than needing to be decided up front.

In this paper we develop a method for inferring the demographic history of a single polyploid population using a diffusion framework that includes homoeologous exchanges and a modified version of the site frequency spectrum (SFS) that collapses data across polyploid subgenomes into a single SFS, removing the need to classify the polyploid as an autopolyploid, allopolyploid, or in between during SNP calling. The collapsed SFS represents the combined sample allele frequencies across the polyploid subgenomes, and is analogous to combining allele frequencies between populations. After describing the model, we compare the SFS generated by this diffusion approximation with frequency spectra generated by forward simulations. We then use forward simulations to assess our ability to infer demographic parameters under various combinations of population bottleneck sizes, bottleneck durations, population divergence times, and homoeologous exchange rates. We then use our model to infer the demographic history of allotetraploid shepherds purse (*Capsella bursa-pastoris*). Finally, we conclude with guidance about the model’s use and interpretation, as well as possible extensions and future directions for work in the area of demographic inference in polyploid species.

## Materials and Methods

### Model description

Our model is inspired by the work of Meirmans and van Tienderen (2013). They focused on the effects of homoeologous exchanges on measures of genetic diversity and population structure in tetraploids by parameterizing an exchange rate, Θ, that determines how frequently alleles are inherited across subgenomes due to homoeologous crossovers and non-disomic inheritance. This exchange rate parameter ranges between 0 and 1, with Θ = 0 corresponding to no allelic exchange and Θ = 1 corresponding to free allelic exchange between subgenomes. Within this framework, the standard categorizations of allopolyploid and tetrasomic autopolyploid fit naturally and represent the extremes in the amount of expected homoeologous crossover. Values of Θ between 0 and 1 allow for intermediate amounts of homoeologous exchange and most closely align with what we would consider to be segmental allopolyploids. We incorporate this range of exchange rates into a diffusion framework, parameterizing homoeologous exchanges akin to migration, much like Roux and Pannell (2015), to provide a generalized model for polyploid demography.

#### A diffusion approximation for polyploids

We considered a single population of a *K*-ploid organism with *S* subgenomes, each containing *k*_1_, *k*_2_, …, *k*_*S*_ chromosomes (∑_*i*_ *k*_*i*_ = *K*). Our diffusion model tracks the joint density of derived mutations across each of the *S* subgenomes, in an infinite sites model. The density of derived mutations at relative frequencies *x*_1_, …, *x*_*S*_ at time *t* is denoted as *φ*(*x*_1_, …, *x*_*S*_, *t*). In general, the relevant diffusion equation is (Kimura 1964)

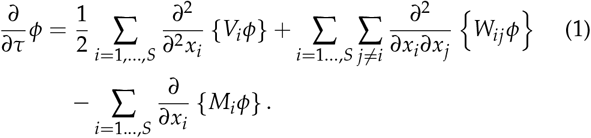

Here, *M*_*i*_ represents the per-generation mean change in the frequency of an allele in subgenome *i, V*_*i*_ the variance in that change, and *W*_*ij*_ the covariance between changes in subgenomes *i* and *j*.

Let *N* be some reference number of individuals (often the ancestral population size) and *ν* be the current relative size of the population. In a population of *N* individuals, there are *Nνk*_*i*_*x*_*i*_ chromosomes in subgenome *i* that carry the derived allele. For a Wright-Fisher model, binomial sampling results in a variance in the number of carriers in the next generation of *Nνk*_*i*_*x*_*i*_ (1 − *x*_*i*_). In general, the binomial distribution has variance *np*(1 − *p*), where *n* is the sample size and *p* is the probability of “success”. In this case, *n* = *Nνk*_*i*_ and *p* = *x*_*i*_, because each of the *Nνk*_*i*_ chromosomes in the next generation is independently copied from a random chromosome in the previous generation, and the proportion of carriers in the previous generation is *x*_*i*_. The allele frequency in the next generation is the number of carriers divided by the total number of chromosomes *Nνk*_*i*_. The variance in the allele frequency in the next generation is thus *x*_*i*_ (1− *x*_*i*_)/*Nνk*_*i*_, because the variance of a random variable *A* divided by a constant *C* is *Var*(*A*)/*C*^2^. Because sampling is independent among subgenomes, all the covariances *W* are 0.

Let *e*_*i↔j*_ be the probability that a meiosis results in an exchange of genetic material between subgenomes *i* and *j*. The expected change in the number of chromosomes in subgenome *I* carrying the derived allele in one generation is then −*Nνe*_*i↔j*_*x*_*i*_ + *Nνe*_*i↔j*_*x*_*j*_. The first term is the loss of alleles due to exchange out of subgenome *i* and the second is the gain due to exchange out of subgenome *j* and into subgenome *i*. The change in the derived allele frequency within subgenome *i* is then

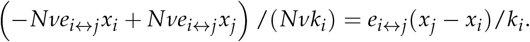

For compatibility with the widely-used diploid equations, we rescaled time to measure in units of 2*N* generations. Defining *E*_*i*_*↔*_*j*_ = 2*Ne*_*i* _*↔*_ *j*_ and Δ*t* = Δ*τ*/(2*N*), plugging in our results for the mean and variance terms, and multiplying both sides of Eq. 1 by 2*N*, we obtained:

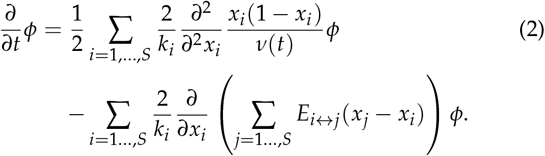

Note that the population size *ν* can be an arbitrary positive function of time, although modelers often employ piecewise constant or exponential functions for *ν*(*t*).

We noted the close analogy between Eq. 2 and the diffusion equation governing alleles within multiple diploid populations (Gutenkunst et al. 2009). The first terms in Eq. 2 model genetic drift and contain an additional constant scaling factor of 2/*k*_*i*_ compared to the typical term. Drift is thus slower in subgenomes with higher ploidy; a tetraploid subgenome (*k*_*i*_ = 4) experiences half the genetic drift of a diploid. This is simply because stochasticity is reduced when there are more chromosomes in the population. This same scaling factor also applies to the rate of mutation influx into each subgenome, compared to the diploid case. The second terms model exchange between subgenomes, which is analogous to migration between populations. The effect on subgenome *i* of exchange with subgenome *j* depends on 2*E*_*i* _*↔*_ *j*_ /*k*_*i*_ compared to the typical migration term *M*_*i* _*←*_*j*_. The additional factor of 2/*k*_*i*_ implies that the effect on the allele frequency within a subgenome of a given influx of alleles decreases as the ploidy increases. Symmetric exchanges between genomes of different ploidy thus have asymmetric effects on allele frequencies. For example, an exchange between a diploid subgenome and a tetraploid subgenome, while biologically rare, could only lead to a 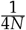 change in allele frequency in a single generation in the tetraploid subgenome versus a potential 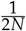 change for the diploid subgenome.

With a distribution for the expected frequency of derived mutations in each subgenome at time *t*, we can generate the expected SFS for a sample of *n* individuals by integrating over the distribution of allele frequencies and calculating the probability of observing *d*_*i*_ derived alleles using a binomial distribution (Sawyer and Hartl 1992):

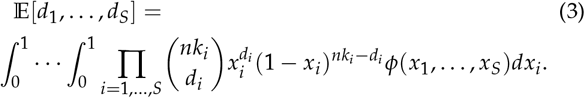

Here each dimension of the SFS corresponds to a different subgenome.

#### Combining polyploid subgenomes

In practice, it may be difficult to partition allele counts between two or more subgenomes. In that case, two or more dimensions of the model SFS must be collapsed down to a single dimension for comparison with the observed SFS. This problem can arise due to issues with phasing, unknown or unsampled parental lineages, or simply unknown origin for the polyploid. As an example, consider a sample of *n* polyploid individuals with two subgenomes with ploidal levels *k*_1_ and *k*_2_. The original SFS for this population, 𝔼 [*d*_1_, *d*_2_],would be a two-dimensional array of size (*nk*_1_ + 1) ×(*nk*_2_ + 1). We collapse the two dimensions together using the following equation:

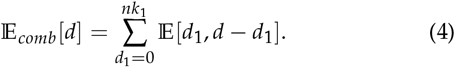

In words, the *d* entry of the reduced SFS is the summation of all entries for which the total allele count in the removed dimensions equals *d*. This collapse of the SFS is illustrated in Fig. 2. The resulting dimension of the new SFS is of size *nk*_1_ + *nk*_2_ + 1 and is what we use for comparison with the observed SFS when performing demographic inference with a polyploid population. If more than two subgenomes are indistinguishable, then this reduction process can be iterated to collapse all indistinguishable genomes into a single dimension of the SFS.

**Figure 1.**
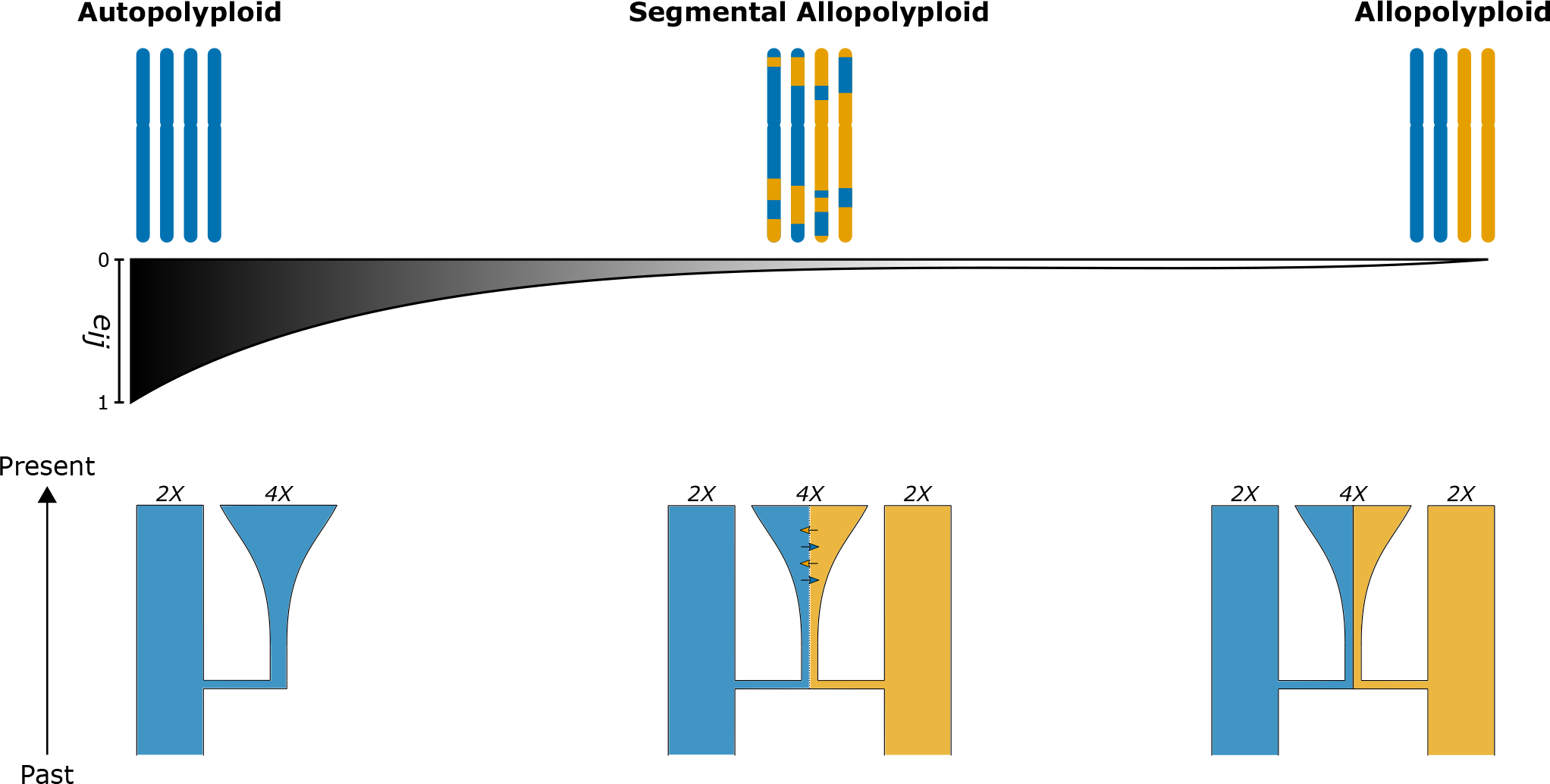
Conceptual representation of the polyploid continuum and corresponding demographic models for polyploid formation. Here *e*_*ij*_ represents the probability of tetrasomic inheritance.

**Figure 2.**
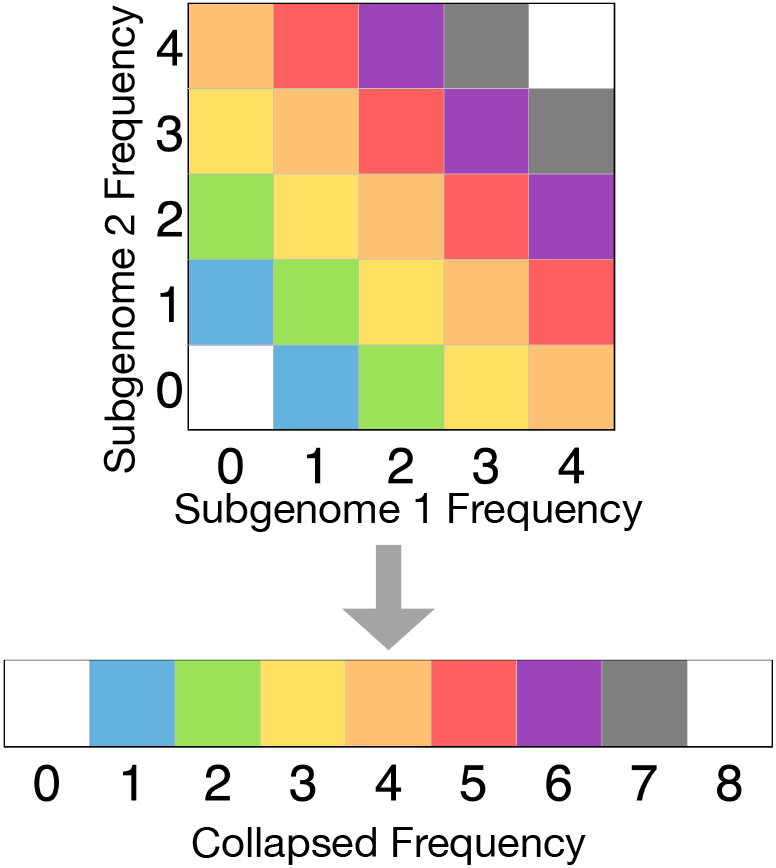
Illustration of SFS collapse across subgenomes. In this case, allele counts and frequencies across two subgenomes are combined, so that the colored entries in the two-dimensional SFS (top) are summed to yield each corresponding entry in the collapsed one-dimensional SFS (bottom).

**Figure 3.**
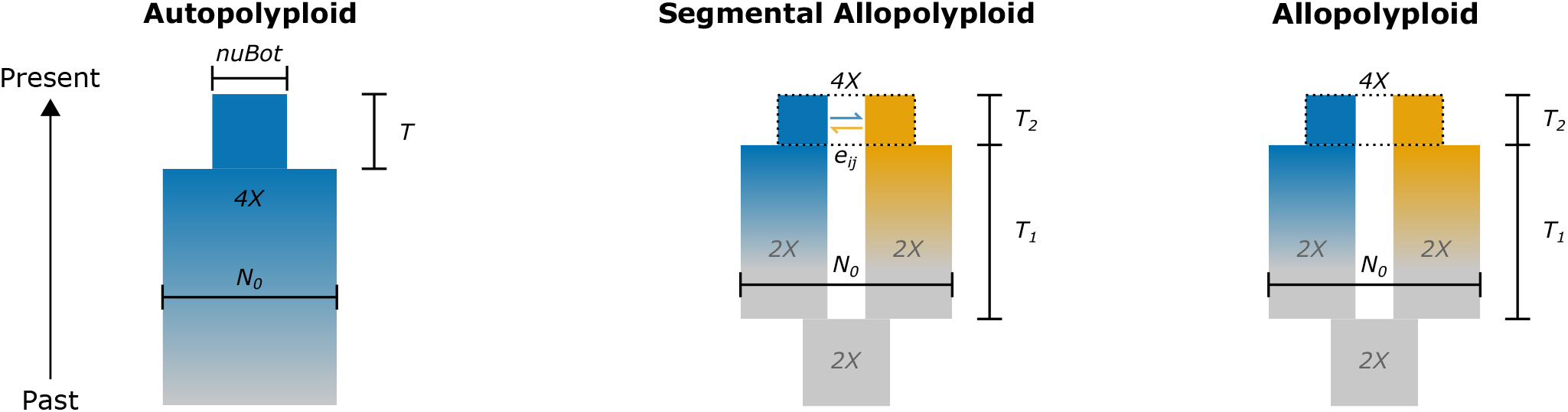
Graphical representations of the models used for validating the diffusion approximation for autopolyploids (left), segmental allopolyploids (middle), and allopolyploids (right). The main parameters used across the models are: *N*_0_ (parental/ancestral population size), *nuBot* (*ν*_*bot*_: proportion of population remaining after bottleneck), *T* (bottleneck duration in autopolyploid model), *T*_1_ (duration of parental divergence before polyploid formation), *T*_2_ (time before sampling for allo- and segmental allopolyploids), and *e*_*ij*_ (*e*_*i*_*↔*_*j*_: per generation probability of homoeologous exchange).

### Validating the diffusion approximation for polyploids

To validate this diffusion approximation, we conducted simulations using SLiM v3.4 (Haller and Messer 2019) under various demographic models and parameter combinations for autotetraploids, allotetraploids, and segmental allotetraploids. For each simulation scenario, we simulated 1,000 polyploid individuals each with a single chromosome 1 Mb in length. Because polyploids have multiple subgenomes, a polyploid individual is composed of multiple SLiM individuals, and therefore subgenomes within individuals were simulated by treating each subgenome as a separate SLiM population. Mutation and recombination rates were set such that *θ* = 2*KNμ* was always equal to 5,000. At the end of each simulation, we generated 50 samples of 10 polyploid individuals by randomly sampling 10 diploid SLiM individuals from each of the SLiM populations and combining them into polyploid individuals to record the SFS. This was repeated 100 times for a grand total of 5,000 simulated frequency spectra for each scenario. We then constructed comparable models using the diffusion approximation implemented in dadi v2.2.0 (Gutenkunst et al. 2009) and compared the expected SFS returned by dadi with the mean SFS across the 5,000 replicates from SLiM to assess how well the two methods corresponded with each other. Specific simulation details for each category of polyploid are given in the following paragraphs.

#### Autopolyploids

For tetrasomic autotetraploids, we constructed a model in SLiM with two populations, representing the two diploid subgenomes that are assumed within the Meirmans and van Tienderen (2013) framework, each containing *N* = 1, 000 diploid SLiM individuals. We set the mutation and recombination rates equal to 6.25 ×10^−7^ and included a symmetric per generation probability of migration equal to 0.5 (*e*_1*↔*2_ = 1.0) to allow free exchange of chromosomes between the subgenomes.

Note that this deviates slightly from our original definition of homoeologous exchanges in that pairs of chromosomes (SLiM individuals), rather than single chromosomes, are moving between subgenomes. We return to this point in the Results and Discussion. An initial burn-in of 40,000 generations was used to reach an approximate state of equilibrium in genetic diversity. The first set of frequency spectra were sampled immediately after this burn-in period to obtain the mean SFS in an equilibrium population. We then simulated bottlenecks of three different sizes (0.2 ×2*N*, 0.5 ×2*N*, and 1.0 ×2*N*), each one lasting for four different lengths of time (1.0 ×2*N*, 2.0 ×2*N*, 3.0 ×2*N*, and 4.0 ×2*N* generations) for a total of 12 parameter combinations.

After simulating data in SLiM, we specified a comparable model in dadi by assuming a single panmictic population with a sample size of 40 chromosomes. Models in dadi start with an equilibrium population, allowing us to immediately generate the expected SFS for a standard neutral model. For the models with bottlenecks, we obtained the expected SFS by including the bottleneck size and duration as parameters to the model and integrating the distribution of allele frequencies forward in time before sampling the resulting frequency spectrum. These expected frequency spectra generated using the diffusion approximation were then compared to the averaged spectra from SLiM to assess their level of agreement by plotting the difference in fit (residuals) between the two methods.

#### Allopolyploids

For fully disomic allotetraploids, we built our model in SLiM with a initial diploid population of 1,000 individuals representing the ancestral reference population. We set the mutation and recombination rates to 1.25 ×10^−6^ and simulated 20,000 burn-in generations to reach approximate equilibrium. After the burn-in period, the ancestral population was split into two populations each containing 1,000 diploid individuals. These separate populations, which represent the ancestors of the allotetraploid subgenomes, were then simulated forward in time at a constant population size for varying numbers of generations (*T*_1_ = 0.5 ×2*N*, 1.0 ×2*N*, 1.5 ×2*N*, and 2.0 ×2*N*) to allow for divergence to develop between the two populations. For the first set of simulations, frequency spectra were sampled immediately after this period of divergence to emulate a newly formed allotetraploid. Within the framework we are proposing here, it is important to note that after this point of polyploid formation the SLiM populations are conceptually a single allotetraploid population. To emulate allopolyploid formation, we also conducted a set of simulations where, after the initial period of divergence between the parents, we introduced bottlenecks of different sizes (*ν* = 0.1 2*N*, 0.25 × 2*N*, and 0.5 × 2*N*), sampling the SFS after differing periods of time (*T*_2_ = 0.25 2*N*, 0.5 × 2*N*, and 1.0 × 2*N* in generations).

The corresponding model specified in dadi began with a single diploid population that was split into two populations.

For the models without bottlenecks, the populations were integrated forwards in time at a constant population size for the same amount of time as in SLiM (*T*_1_) before sampling the twodimensional SFS (20 chromosomes per SLiM population) and combining it into a one-dimensional frequency spectrum using Eq. 4. For models with bottlenecks, the two populations were once again integrated forwards at a constant size for time *T*_1_ before experiencing an instantaneous bottleneck lasting for time *T*_2_. Frequency spectra from these populations were sampled and combined in the same way and compared to the frequency spectra from SLiM.

#### Segmental allopolyploids

For segmental allotetraploids, the simulation setup was similar to the one used for allotetraploids. We included the same initial period of divergence (*T*_1_), as well as the secondary period of time after polyploid formation (*T*_2_). During this secondary period, we added two levels of allelic exchange between subgenomes. The levels we chose were *ei*_*↔*_ *j* = 5 ×10^−5^ and *e*_*i* _*↔*_*j*_ = 5 ×10^−7^. These levels correspond to one exchange event every 10 or every 1,000 generations, respectively. Parameters for the initial period of divergence, *T*_1_, were kept the same. We also did simulations with bottlenecks, using bottlenecks of the same sizes, *ν*, during the time period *T*_2_ for the corresponding set of models. Setting up the model in dadi for segmental allotetraploids was identical to allotetraploids except with the addition of the exchange parameter, *E*_*i* _*↔*_ *j*_ = 2*Ne*_*i* _*↔*_ *j*_, during the integration for *T*_2_.

### Parameter inference and identifiability

In a separate set of SLiM simulations, we also sought to understand how well demographic parameters could be inferred from a combined polyploid SFS in dadi. For these simulations, we used a subset of the parameterizations listed above for allotetraploids and segmental allotetraploids, simulating 10 independent frequency spectra for each parameter combination. We also included an additional layer of complexity in these simulations by incorporating two different types of data generation meant to mimic data collected using genotyping by sequencing (GBS) and whole-genome resequencing data (WGS). For each GBS simulation, we generated 100 bp segments across 5000 independent SLiM runs and combined them into a single SFS. For each WGS simulation, we generated 5 Mb regions across 10 independent SLiM runs and combined them into a single SFS.

Models in dadi were specified as described in the previous section and were used to maximize the composite likelihood for the simulated input data. Parameters were initialized at random starting points and were estimated using the nlopt-enabled optimizer in dadi (BOBYQA algorithm; Powell 2009; Johnson 2014). We conducted 50 independent optimization replicates for each simulated data set for all models and simulation parameters to assess convergence on the same set of maximum likelihood parameter estimates from different starting points. After this, we used R v3.6 to sort the results by likelihood value, keeping the parameters producing the highest likelihood from the 50 optimizations across the twenty replicates for each simulation, and compared these estimates to the true value used to simulate the data.

### Empirical example: *Capsella bursa-pastoris*

To model the demographic history of shepherd’s purse, we first obtained the multidimensional SFS with the *C. bursa-pastoris* subgenomes separated from Douglas *et al*. (2015). We then combined the subgenomes into a one-dimensional frequency spectrum with Eq. 4 and used this SFS as input for parameter inference in dadi. Within dadi, we specified two models for comparison: the allotetraploid bottleneck model and the segmental allotetraploid bottleneck model. These two models are identical apart from the inclusion of allelic exchange between subgenomes in the segmental allotetraploid version. Parameters for each model were estimated with the BOBYQA algorithm in dadi using 100 independent optimization runs from different random starting points. We then used the parameter estimates with the best likelihood to compare the models with the observed data, as well as conducting a likelihood ratio test. Confidence intervals were estimated at the 95% level assuming unlinked sites using the Fisher Information Matrix (Coffman *et al*. 2015) and propagation of uncertainty for composite parameters (population sizes, times, and migration rates) as described in Blischak *et al*. (2020). Parameters were converted from dadi units to real units using a mutation rate of 7 ×10^−9^ and total sequence length (*L*) of 773,748 bp, the same values used by Douglas *et al*. (2015). As a secondary comparison, we also recreated the best-fitting model for the *C. bursa-pastoris* subgenomes from Douglas *et al*. (2015, model C), which included exponential growth in the popular tions after formation, and compared the results to those from our models.

### Data availability

Supplemental files, including scripts for performing simulations, analyzing results, and generating plots, can be found on GitHub (https://github.com/pblischak/polyploid-demography.git) and are archived on figshare (https://doi.org/10.6084/m9.figshare.20635635.v3). All simulated data files, optimization results, and data files for analyses of *C. bursa-pastoris*, are also available on both GitHub and figshare.

## Results and Discussion

We developed a generalized diffusion approximation for polyploids by extending previous work on the multi-population diffusion equation. Within this framework, we are able to accommodate the full continuum of polyploid formation types by explicitly parameterizing homoeologous exchanges between subgenomes. We then used simulations to validate the diffusion approximation by comparing it to results from forward simulations using collapsed frequency spectra to emulate the difficulties of separating parental subgenomes. We also investigated the accuracy of parameter inference under the diffusion framework for a subset of parameter values for allotetraploids and segmental allotetraploids. Finally, we used the polyploid diffusion model to infer the demographic history of *Capsella bursa-pastoris*, a widely distributed allotetraploid, finding evidence for allelic exchange between subgenomes.

### The diffusion approximation in polyploids

We compared the expected SFS from the polyploid diffusion approximation as implemented in dadi with frequency spectra generated by forward simulations in SLiM. Figure 4 shows these results for a sample of parameter combinations for models including bottlenecks across autotetraploids, allotetraploids, and segmental allotetraploids. For these examples, as well as for the other parameters used for the simulations, we find good qualitative agreement between the frequency spectra from dadi and SLiM.

**Figure 4.**
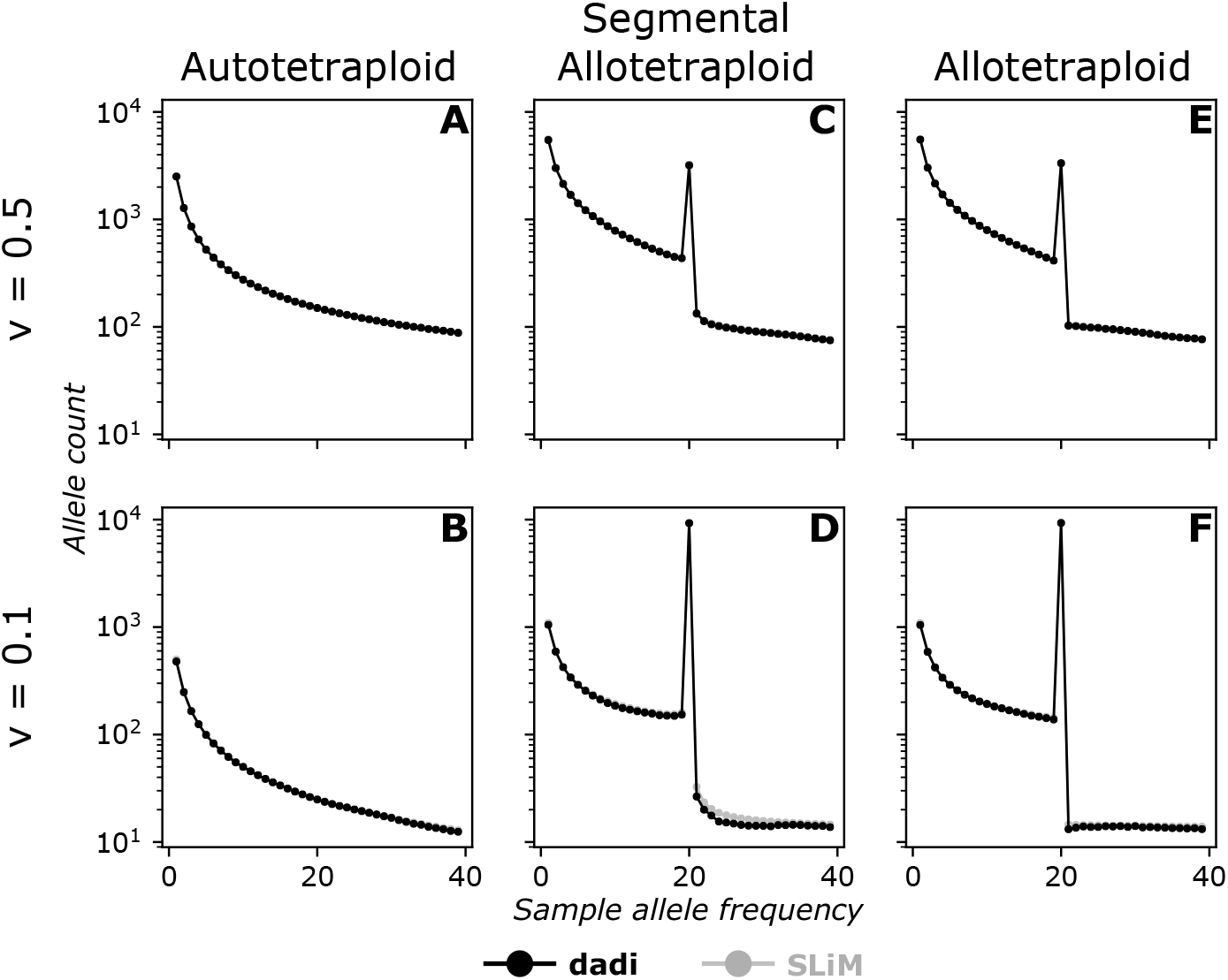
Comparison of collapsed frequency spectra between SLiM (black) and dadi (grey) for two different bottleneck sizes (*ν* = 0.5 [top row] and *ν* = 0.1 [bottom row]) for autotetraploids (A & B), segmental allotetraploids (C & D), and allotetraploids E & F). The rate of homoeologous exchange for segmental allotetraploids was set to *e*_*i↔j*_ = 5 ×10^−6^.

As might be expected for autopolyploids, the frequency spectra appear similar to what we would obtain for a diploid but with double the sample size (Fig. 4A&B). This is the case even though we simulated the autotetraploid in SLiM as two populations forming the subgenomes of a single polyploid population. For allotetraploids and segmental allotetraploids, combining allele frequencies between divergent parental lineages results in a characteristic spike in sites with allele frequencies of 50% (Fig. 4C-F). Much of this pattern is driven by opposite alleles drifting toward fixation in the two subgenomes, leading to fixed heterozygosity. This is more pronounced in the frequency spectra for populations experiencing a stronger bottleneck (*ν* = 0.1 versus *ν* = 0.5), where the spike at 50% frequency is higher and the drop off in the prevalence of sites with allele frequencies over 0.5 is greater. Segmental allotetraploids also differ in the appearance of their 50% frequency spike, having small shoulders of increased counts of sites with frequencies around 50% (Fig. 4C&D). This is caused by the allelic exchange between subgenomes generating allelic combinations that are not possible in allotetraploids due to the complete separation of subgenomes. As the exchange rate in segmental allotetraploids increases, the spike continues to level out and eventually becomes visually indistinguishable from an autotetraploid (Fig. 5).

**Figure 5.**
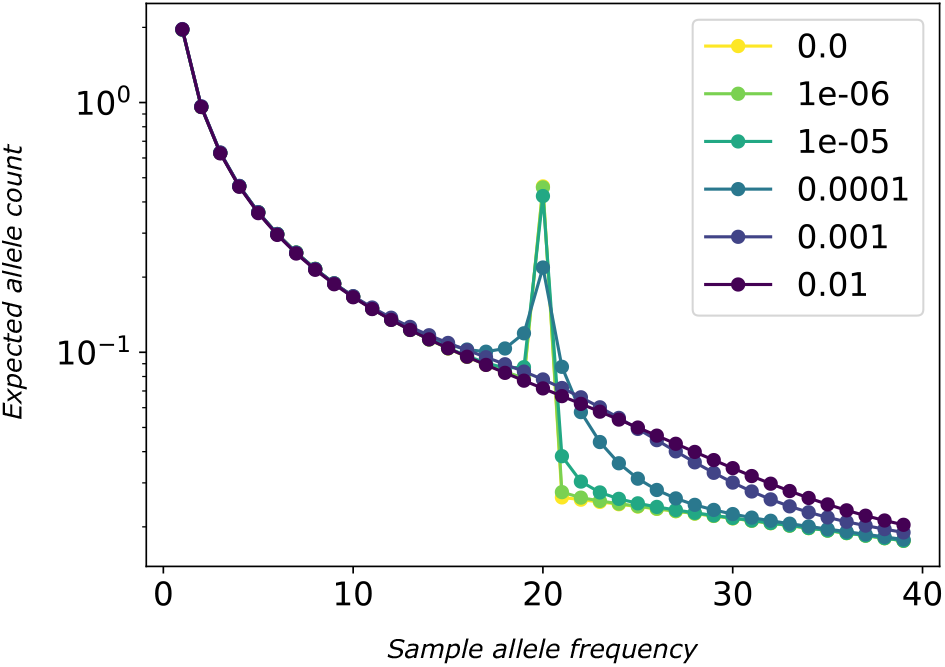
Illustration of the effect of the homoeologous exchange rate on the collapsed polyploid site frequency spectrum for a tetraploid. Here exchange rates vary from *e*_*i*_*↔*_*j*_ = 0 (allotetraploid) to *e*_*i*_*↔*_*j*_ = 0.01. At the high end of this range, the SFS no longer has a distinctive peak at 50% frequency due to the exchanges mixing alleles, making the SFS appear more similar to that of an autopolyploid.

### Inferring demographic parameters in polyploids

As a follow-up to our validating simulations, we also sought to understand how well we could infer demographic parameters using the numerical approaches implemented in dadi for collapsed allotetraploid and segmental allotetraploid frequency spectra. For the parameters that describe the formation of the polyploid population itself, we are typically able to obtain precise parameter estimates across our simulated scenarios (Fig. 6). As expected, parameter estimates for the GBS data simulations generally show more variation than the WGS simulations. However, the WGS simulations include linkage, and demonstrate the accuracy of parameter inference even when the assumption of independent sites is violated.

**Figure 6.**
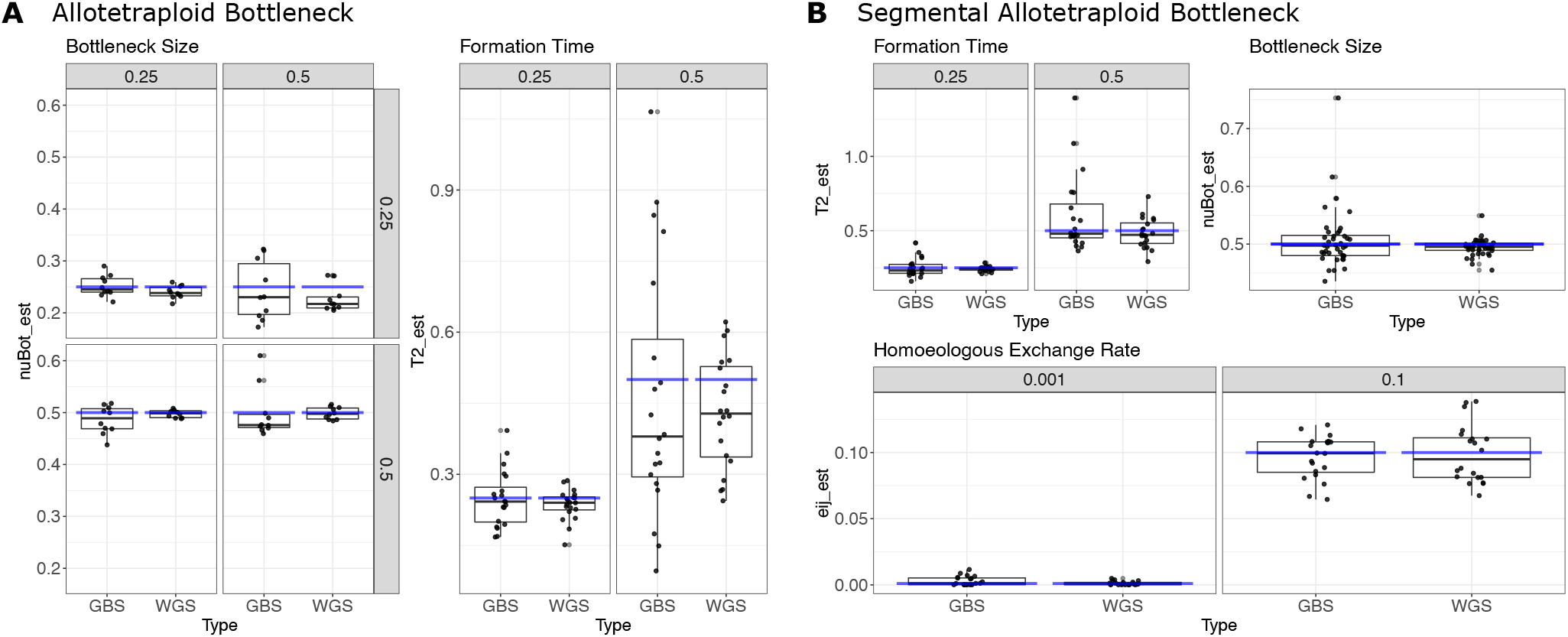
(A) Parameters estimates from dadi for bottleneck size (left panel) and formation time (right panel) for the allotetraploid bottleneck model simulated with SLiM across two different data types: Genotyping-by-sequencing (GBS) and whole-genome resequencing (WGS). For estimates of the bottleneck size, the secondary divisions in the plots show the true formation time (*T*_2_ = 0.25, 0.5) in the rows and the true bottleneck size (*ν*_*Bot*_ = 0.25, 0.5) in the columns. (B) Parameters estimates from dadi for formation time (top-left panel), bottleneck size (top-right panel), and homoeologous exchange rate (bottom panel) for the segmental allotetraploid bottleneck model simulated with SLiM across GBS and WGS data types. For all plots, the blue line represents the true value used to simulate the data.

Estimates for parameters that describe the demographic model before the formation of the polyploid are generally less precise and often showed unstable behaviors when searching for optimal values (see Supplemental Files). For example, estimates of the combined parental population sizes (denoted *N*_0_ in our models) were consistently unstable/unbounded. This suggests that parameters estimated for the parental lineages of polyploid populations from a one-dimensional collapsed frequency spectrum should be interpreted with caution. For diploid populations and piecewise constant population histories, such instabilities have been well characterized by the geometry of the SFS and the effect of sampling error in the SFS on estimating parameter values on the boundary of the search space (Rosen *et al*. 2018).

### Demographic history of *Capsella bursa-pastoris*

Shepherd’s purse (*Capsella bursa-pastoris*; Brassicaceae) is a well-studied allotetraploid species resulting from hybridization between an outcrossing species, *C. grandiflora*, and a selfing species, *C. orientalis*. Previous work on the demographic history of *C. bursa-pastoris* found that it formed roughly 100 kya, with a current distribution spreading across Europe, the Middle East, and Asia (Douglas *et al*. 2015; Roux and Pannell 2015; Kryvokhyzha *et al*. 2019). These three regions also correspond to three major genetic groups within the species, all of which have experienced varied evolutionary trajectories including bottlenecks and changes in life history, though the Middle Eastern region is where the species is inferred to have originated (Cornille *et al*. 2016). Here we use data for shepherd’s purse from Douglas *et al*. (2015) as an empirical example to compare the results of conducting demographic inference with a one-dimensional collapsed SFS to those obtained by the original study.

Using a collapsed representation of the SFS from Douglas *et al*. (2015), we modeled the demographic history of *Capsella bursa-pastoris* using the allotetraploid bottleneck and segmental allotetraploid bottleneck models (Table 1). Both models resulted in similar estimates for the parameters regarding the formation of the polyploid lineage (*ν*_*bot*_ and *T*_2_; see Fig. 3), finding that *C. bursa-pastoris* was formed around ∼ 270–285 kya with an initial effective population size of ∼ 50,000–55,000 individuals. These estimates differ somewhat from the estimates reported in (Douglas *et al*. 2015), who found that *C. bursa-pastoris* formed more recently, between ∼22–177 kya. However, Douglas *et al*. (2015) estimated separate effective populations sizes for each subgenome, finding similar values to our combined estimate: ∼ 6,000–55,000 individuals for the *C. grandiflora* subgenome (*C. bursa-pastoris* A) and ∼ 12,000–101,000 for the *C. orientalis* subgenome (*C. bursapastoris* B). For most other parameters, the allotetraploid and segmental allotetraploid models had drastically different esti-mates of the ancestral and combined parental population sizes (*N*_*A*_ and *N*_0_, respectively) and a roughly 0.75X difference in the estimate of parental divergence (*T*_1_). This is consistent with the instability observed in our simulation studies for pre-polyploid formation parameters. The estimated exchange rate (*e*_*i↔j*_) for the segmental allotetraploid model was 6 ×10^−8^, suggesting that rare bouts of allelic exchange may have occurred between the subgenomes of *C. bursa-pastoris*. This is corroborated by the fact that the composite log-likelihood for the segmental allotetraploid model was *∼* 138 units higher than for the allotetraploid model, resulting in a likelihood ratio test statistic of 277, corresponding to a vanishingly small p-value.

**Table 1.** Parameter estimates for *Capsella bursa-pastoris* demographic history. Population sizes (*N*_*\**_) are reported as the number of individuals and time intervals (*T*_*\**_) are reported as the number of years. Ranges for 95% confidence intervals are given in parentheses.

| Parameter | Allotetraploid Bottleneck | Segmental Allotetraploid Bottleneck |
| --- | --- | --- |
| $N_A$ | 249,000 (210,000 – 296,000) | 7,720 (4,830 – 12,400) |
| $N_0$ | 24,900,000 (15,800,000 – 39,300,000) | 772,000 (555,000 – 1,070,000) |
| $N_{bot}$ | 50,400 (39,000 – 65,100) | 55,300 (39,900 – 76,800) |
| $T_1$ | 1,030,000 (792,000 – 1,330,000) | 1,540,000 (1,110,000 – 2,150,000) |
| $T_2$ | 269,000 (208,000 – 347,000) | 284,000 (204,000 – 394,000) |
| $e_{i \leftrightarrow j}$ | – | $6.0 \times 10^{-8}$ ($4.4 \times 10^{-8}$ – $8.2 \times 10^{-8}$) |
| % misidentified | 1.66 (1.40 – 1.97) | 3.14 (1.96 – 5.02) |

Our estimates of the timing and population size impacts of polyploid formation for *C. bursa-pastoris* are similar to those of Douglas *et al*. (2015), but our estimates of pre-polyploid parameters differ substantially. This is not unexpected, given that our simulation experiments exhibited unbounded behavior in likelihood optimization for those parameters. Furthermore, the parental taxa forming *C. bursa-pastoris* have different life history strategies, with *C. orientalis* being highly inbred. This could be leading to the differences we see in our estimates of divergence between the parental lineages and their effective population sizes. Incorporating this inbreeding into the model should be possible using sampling schemes that build inbreeding in their derivation of the expected SFS (Blischak *et al*. 2020).

Another important distinction between our analyses and those of Douglas *et al*. (2015) is their use of a four-population model for their demographic inferences, including both parents, *C. grandiflora* and *C. orientalis*, and separating the corresponding subgenomes of *C. bursa-pastoris*. In our combined analysis, we are deliberately excluding this additional information to better understand the limits of demographic inference within a single polyploid population. The result is that we are not able to reliably estimate all model parameters. However, when we do compare our models with a collapsed version of the SFS generated by recreating the Douglas *et al*. (2015) model, we see that the Douglas *et al*. (2015) model more closely resembles the allotetraploid bottleneck model (Figure 7). Furthermore, the log-likelihood for the reproduced Douglas *et al*. (2015) model is -789.14, which is 276/ log-units lower than the log-likelihood for our segmental allotetraploid model (−512.91). And although we have focused on collapsed spectra, generating 2D spectra to match the separated subgenomes in the original SFS shows that including exchange, even though there appears to be no shared variation in the 2D SFS for the data, also improves the log-likelihood by ∼ 230 units (−1270.60 for the 2D segmental allotetraploid model vs. -1501.87 for the 2D Douglas *et al*. (2015) model; Supplemental Figure S1). It is unclear why this discrepancy exists between the models and why a model predicting exchange provides a better fit even though the data shows no shared variation, but it could suggest the occurrence of rare homoeologous exchanges between the two subgenomes of shepherd’s purse. Future work using our modeling framework with the inclusion of parental lineages, inbreeding, and other demographic factors will help us to better disentangle the evolutionary forces affecting *C. bursa-pastoris* as well as other polyploid species.

**Figure 7.**
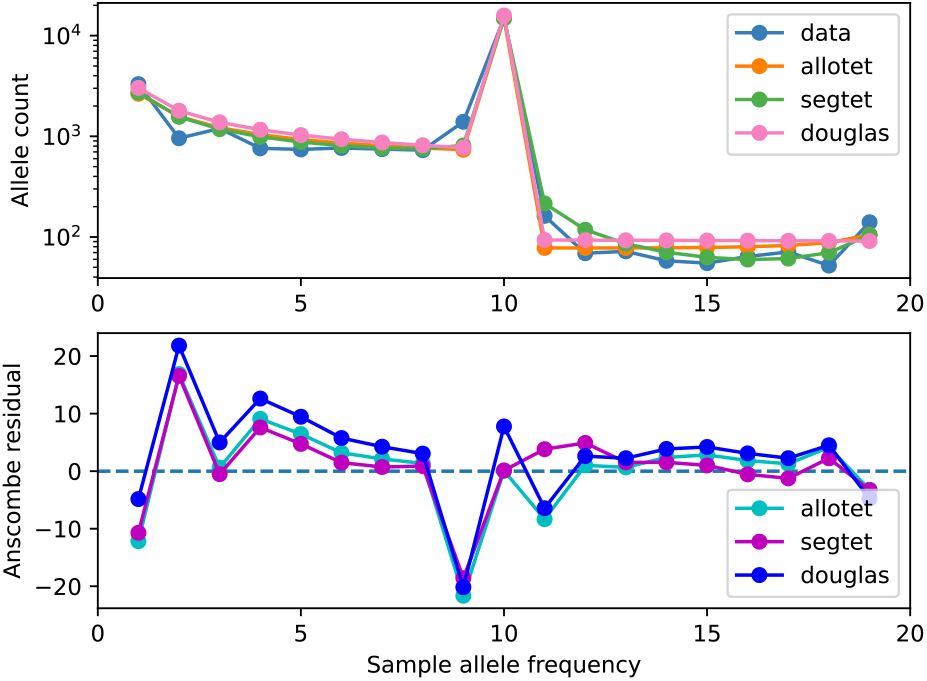
[top] Site frequency spectra resulting from the maximum likelihood parameters estimated for the allotetraploid bottleneck and segmental allotetraploid bottleneck models for *C. bursa-pastoris*, as well as for the exponential growth model from Douglas *et al*. (2015). The observed data are also shown in blue. [bottom] Anscombe residual plot (model - data) comparing each entry in the SFS between the allotetraploid bottleneck, segmental allotetraploid bottleneck, and Douglas *et al*. (2015) models with the observed data.

### General considerations for modeling the demographic history of polyploids

Researchers studying polyploid taxa are faced with numerous challenges when analyzing genomic data to understand more about the evolutionary history of their study organism(s). Here we have proposed a model to help alleviate some of these issues by explicitly parameterizing the continuum of polyploid formation types based on the expectations for a collapsed polyploid SFS. Given the results of our simulation and empirical analyses, there are clear advantages and disadvantages to an-alyzing data in this way. One advantage is that there is no longer a need to separate variation occurring within potential subgenomes, because the model accommodates auto-, allo-, and segmental allopolyploids. Assuming complete separation of subgenomes in an allopolyploid or completely free recombination in an autopolyploid can lead to unintentionally ignoring signals for intermediate patterns that may have important consequences. Furthermore, there may be only certain parts of the genome that experience exchanges. For example, allotetraploid *Arabidopsis suecica*, while primarily having bivalent pairing of chromosomes within each of its subgenomes, was found to have variation in patterns of gene expression owing to a relatively small number of homoeologous exchanges present in some samples (Burns et al. 2021). In this case, the authors were able to leverage the robust genomic resources available in *Arabidopsis* to recognize a signal for homoeologous exchange. For most species, however, identifying the mode of formation will have to be done in the absence of a reference genome or genomes. With our model, the shape of the collapsed SFS can be a good initial indicator of whether the species is an autoor allopolyploid, with shoulders around any peaks at 50% frequency providing additional evidence for the presence of homoeologous exchanges. Further investigation using the likelihood-based framework in dadi to perform model comparisons for determining the mode of formation could also provide a robust means of identifying even small amounts of homoeologous exchange in polyploid lineages.

Several other nuances in the demography of polyploids that warrant further investigation within the framework we’ve proposed include modeling biases in the genomic regions experiencing homoeologous exchanges, the process of diploidization and the shift from tetrasomic to disomic inheritance (particularly in autopolyploids), and distinguishing between homoeologous exchanges and the retention of ancestral polymorphism (incomplete lineage sorting; ILS). For biases in homoeologous exchange, previous work on barriers to gene flow provide a compelling starting point. For example, using a similar setup to Tine *et al*. (2014), who used dadi to investigate non-uniform patterns of gene flow within the genomes of European sea bass, we performed a small simulation to investigate restricting homoeologous exchanges to only a certain proportion of the genome, finding that these biases do impact the shape of the SFS (Supplemental Figure S2). Capturing the decay of tetrasomic inheritance could be possible by making the homoeologous exchange rate time dependent and using a demographic model akin to the isolation-with-initial-migration model (Wilkinson-Herbots 2012), allowing for the estimation of the onset of disomic inheritance. Distinguishing between homoeologous exchange and ILS would involve similar analyses involving isolation-with-migration (Nielson and Wakeley 2001), since homoeologous exchange is similar to gene flow from a modeling perspective.

The primary disadvantage of analyzing data using a collapsed spectrum that combines allele frequencies across subgenomes is the inability to reliably infer ancestral dynamics prior to polyploid formation. This pattern was observed across our simulations and the analysis of *C. bursa-pastoris*. As we mentioned above, one way to potentially deal with these issues would be to include the parental lineages in the demographic model. This does require additional knowledge about the study system but should be feasible within dadi, if the data are available, by adding up to four or five populations using the newly implemented graphics processing unit acceleration (Gutenkunst 2021). Theoretical investigations into the limitations of using a collapsed polyploid frequency spectrum could also further illuminate parameter and model identifiability and could be used to guide additional innovation in the construction of more informative demographic models for polyploids.

## Conclusions

Disentangling the roles of demography and selection in polyploids will be a major step toward better understanding the role they play in the generation and maintenance of biodiversity. Additionally, an appreciation for the continuum-like nature of polyploids will remain essential as more methods are developed to model their evolutionary histories. The method we have developed here provides a foundation for further exploration of diffusion-based demographic models for polyploids and reveals important pain points and considerations for how to approach demographic modeling with a collapsed polyploid SFS. Including parental lineages and gaining a better theoretical understanding of the effect of fixed differences between subgenomes on demographic inference will be key advancements in future iterations of our modeling approach. Combining more robust demographic models for polyploids with existing frameworks for studying selection (e.g., Huang *et al*. 2021) will then provide a powerful framework for revealing their evolutionary importance in the short term.

## Acknowledgements

The authors thank Justin Conover, Camille Roux, and two anonymous reviewers for comments that helped to improve the manuscript, and Stephen Wright for sharing the data for *Capsella bursa-pastoris*. We also would like to thank J. Wendel, T. Steussey, and attendees of the Polyploid Webinar (https://www.barkerlab.net/polyweb) for their thoughtful input regarding effective population sizes in polyploids, the nature of segmental allopolyploidy, and for exhibiting a general enthusiasm for all things whole-genome duplication.

