## Supplemental Figures for "Demographic History Inference and the Polyploid Continuum"

#### SUPPLEMENTAL MATERIALS

Paul D. Blischak, Mathews Sajan, Michael S. Barker, and Ryan N. Gutenkunst

#### List of Figures

S1 2D spectra comparing our model with Douglas *et al.* . . . . . . . . . . . . . . . . . . . . . . . . . . . 2
S2 SFS for non-uniform homoeologous exchanges across the genome . . . . . . . . . . . . . . . . . . 3

### Supplemental Figures

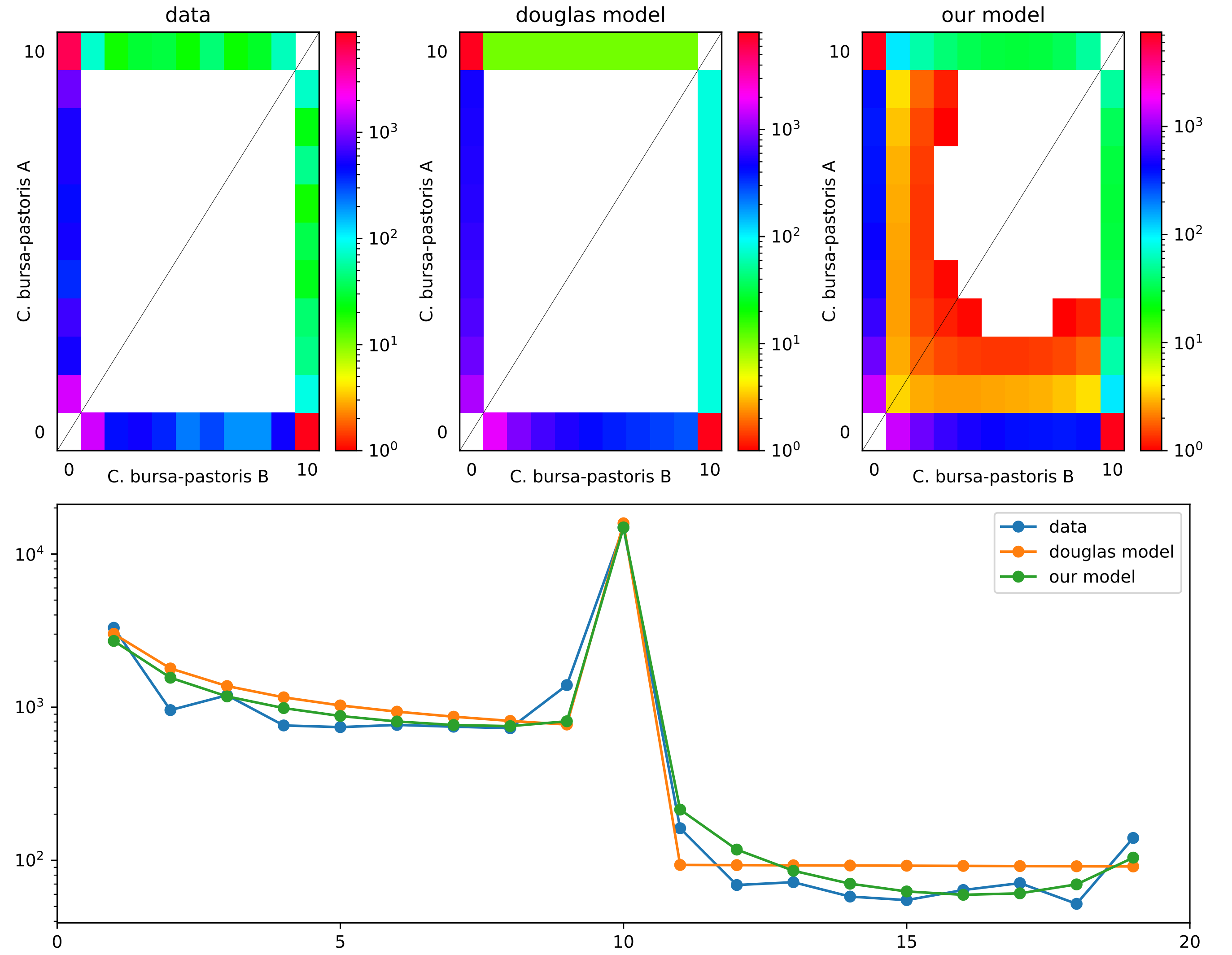


**Fig. S1.** [top row] 2D plots of the site frequency spectrum for the A and B subgenomes of *Capsella bursa-pastoris* used in Douglas *et al.* (2015), as well as the SFS produced by their model with exponential growth and the SFS from our segmental allotetraploid model, from left to right respectively. [bottom row] Collapsed, 1D version of the SFS corresponding to the 2D versions in the top row.

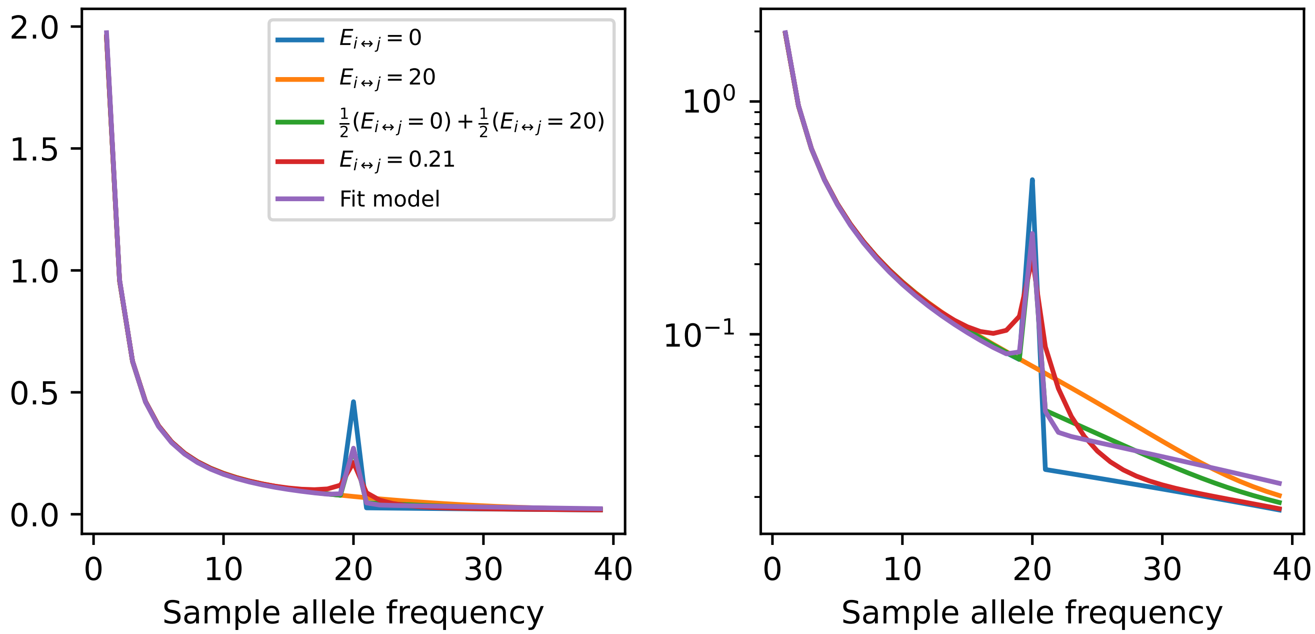


**Fig. S2.** Here we show five spectra that illustrate different properties associated with the amount and type of homomoeologous exchanges that may be experienced by a polyploid lineage. The blue spectrum ($E_{i\leftrightarrow j} = 0$), corresponds with a typical allopolyploid model where no exchange is happening between subgenomes. The orange spectrum ($E_{i\leftrightarrow j} = 20$), corresponds to extremely high levels of homoeologous exchange such as that of a tetrasomc autopolyploid. The green spectrum ($\frac{1}{2}E_{i\leftrightarrow j} = 0 + \frac{1}{2}E_{i\leftrightarrow j} = 20$), is a mixture of the previous two spectra and represents a pattern of homoeologous exchanges that are non-uniform across the genome. The red spectrum ($E_{i\leftrightarrow j} = 0.21$), shows the pattern of homoeologous exchange that we associate with segmental allotetraploidy and is random across the genome. The fifth spectrum, in purple, shows the result of fitting a segmental allotetraploid model (random exchange) to the non-uniform exchange model. From these spectra, we can see that non-uniform exchanges show different patterns in the SFS than when exchanges are modeled as random, which could lead to misinterpretation of the mode of formation of the polyploid lineage and incorrectly inferred parameter values.
